# Fiber tractography using machine learning

**DOI:** 10.1101/104190

**Authors:** Peter F. Neher, Marc-Alexandre Côté, Jean-Christophe Houde, Maxime Descoteaux, Klaus H. Maier-Hein

## Abstract

We present a fiber tractography approach based on a random forest classification and voting process, guiding each step of the streamline progression by directly processing raw diffusion-weighted signal intensities. For comparison to the state-of-the-art, i.e. tractography pipelines that rely on mathematical modeling, we performed a quantitative and qualitative evaluation with multiple phantom and *in vivo* experiments, including a comparison to the 96 submissions of the ISMRM tractography challenge 2015. The results demonstrate the vast potential of machine learning for fiber tractography.

## 1. Introduction

Fiber tractography on the basis of diffusion-weighted magnetic resonance imaging (DW-MRI) has been a research topic for almost 20 years. A vast spectrum of tractography algorithms has been presented over the last years ranging from local deterministic approaches (Basser, 1998; Chao et al., 2008; Lazar et al., 2003; Mori et al., 1999; Tournier et al., 2012) through probabilistic methods (Behrens et al., 2007; Berman et al., 2008; Descoteaux et al., 2009; Friman et al., 2006; Vorburger et al., 2013; Zhang et al., 2013) to global tractography (Aganj et al., 2011; Daducci et al., 2015; Fillard et al., 2009; Jbabdi et al., 2007; Lemkaddem et al., 2014; Mangin et al., 2013; Reisert et al., 2011). To infer information about the complex microstructure of brain tissue and to optimally exploit the acquisition dependent signal characteristics of diffusion-weighted images, tractography algorithms employ mathematical models calculated from the diffusion-weighted signal. Prominent examples include the diffusion tensor (Basser et al., 1994), multi tensor models (Kreher et al., 2005; Malcolm et al., 2010), spherical deconvolution (Alexander, 2005; Jeurissen et al., 2014; Schultz et al., 2010; Tournier et al., 2007), persistent angular structures (Jansons and Alexander, 2003), Q-ball modelling (Aganj et al., 2009; Descoteaux et al., 2007) as well as a large variety of multi-compartment models (Assaf et al., 2008; Assaf and Basser, 2005; Panagiotaki et al., 2012; Sotiropoulos et al., 2012; Zhang et al., 2012). To obtain such a model representation of certain local tissue properties, the corresponding inverse problem has to be solved using the measured signal. Depending on the model, this imposes different constraints on the data quality and acquisition sequence, such as a minimum number of diffusion-weighting gradients and a high signal to noise ratio (SNR). Usually, the more expressive a model is, the more demanding is its calculation and the higher are the requirements for the signal. Choosing the optimal model is not a trivial problem. Oversimplified modelling can, for example, hamper the ability to resolve crossing fiber situations. Modeling approaches make various assumptions about signal and tissue properties that are highly variable across subjects, locations in the brain and acquisition schemes. This issue has been discussed extensively and there is still no solution that is optimal in all situations (Daducci et al., 2014; Farquharson et al., 2013; Jbabdi and Johansen-Berg, 2011; Neher et al., 2015a; Nimsky, 2014). Furthermore, depending on the model and the dataset, a certain number of tractography parameters, such as a termination criterion, e.g. on the basis of a threshold on the fractional anisotropy (FA), have to be adjusted manually, which requires expert knowledge.

In the context of signal modeling, initial studies have successfully shown the potential of machine learning techniques, e.g. for the tasks of image quality transfer and tissue micro-structure analysis (Alexander et al., 2014; Golkov et al., 2016; Nedjati-Gilani et al., 2014; Reisert et al., 2016) and to estimate the number of distinct fiber clusters per voxel (Schultz, 2012). These methods avoid or alleviate some of the issues that come with the usage of diffusion-signal models.

Here, we present the first approach to fiber tractography on the basis of machine learning, which has several advantages: There is no need to explicitly solve the inverse problem to obtain a representation of the diffusion propagator or the tissue microstructure from the diffusion-weighted signal. Also, classical modeling approaches often struggle with artifacts that are not included in the mathematical model, such as noise and distortions, while a machine learning based approach can, to a certain extent, deal with such signal imperfections by learning them from the training data. While the desired information has of course to be encoded in the signal, there are no general restrictions on the type of image acquisition, e.g. regarding the number of diffusion-weighting gradients or the b-value. This enables straight-forward optimization of the method to a specific acquisition scheme. Furthermore, the distinction between white matter and non-white matter tissue is directly learned from the training data. This means that additional white matter mask images or model derived thresholds, such as on the FA or on the magnitude of the peaks determined by the model, are not necessary to constrain the tractography. Additionally, in classical streamline tractography, the decision about the next direction of the streamline progression is typically based solely on the signal information at the current streamline position. The probabilistic nature of our approach enables the meaningful integration of the information obtained from multiple signal samples in the local neighborhood, guiding each step of the streamline progression.

This work is based on the preliminary results and methods presented at MICCAI 2015 (Neher et al., 2015b). Here, we introduce new types of classification features and training data. The evaluation of our method was extended to the data and results from the ISMRM tractography challenge 2015, including 96 tractography methods for comparison. Furthermore, we extensively assessed the capabilities of our method to generalize from *in vivo* to *in vivo*, *in vivo* to *in silico* and *in silico* to *in vivo* data. In our experiments, we show that the method performs as good as or better than the state-of-the-art and generalizes well to unseen datasets.

## 2. Materials and Methods

Standard streamline tractography approaches reconstruct a fiber by iteratively extending the fiber in a direction depending on the current position. The directional information is usually inferred from a signal model at the respective location, such as the diffusion tensor (DT), fiber orientation distribution functions (fODF) or diffusion orientation distribution functions (dODF). The method presented in this work also iteratively extends the current fiber but in contrast to standard streamline approaches, the determination of the next progression direction relies on a different concept:

1. Instead of mathematically modeling the signal the presented method employs a random forest classifier working on the raw diffusion-weighted image values in order to obtain information about local tissue properties, such as tissue type (white matter or not white-matter) and fiber direction (cf. sec. 2.1).
2. To progress or possibly terminate a streamline, not only the image information at the current location but also at several sampling points distributed in the neighborhood are taken into account. The final decision on the next action is then based on a voting process among the individual classification results at these sampling points (cf. sec. 2.2).

### 2.1. Learning fiber directions using random forest classification

#### Classification features

We evaluated two types of input features for the random forest classifier: raw signal intensities as well as the voxel-wise coefficients of the corresponding spherical harmonics fit of the diffusion-weighted signal. In both cases, multiple b-values can be handled by concatenating the feature vectors of the individual b-shells. For the first case, the signal is resampled to 100 directions equally distributed over the hemisphere using spherical harmonics, to become independent of the gradient scheme used to acquire the data. In addition to the diffusion-weighted signal features, the normalized previous streamline direction is added to the list of classification features, thus enabling the method to better overcome ambiguous situations. Each signal feature is used twice for training, one time in conjunction with the directional feature and a second time with a zero-vector instead of the direction feature to enable valid classifications at streamline seed points. We furthermore investigated the effect of adding features such as *T1* signal values or scalar indices derived from the diffusion-weighted signal, such as the generalized fractional anisotropy (GFA) (Tuch, 2004).

#### Reference directions

To train the classifier, reference fiber tracts corresponding to the respective diffusion-weighted image are necessary. In our experiments, we explore two variants of obtaining these reference tracts. (1) we use a previously performed standard tractography to obtain the reference tracts. The impact of the tractography algorithm choice on the presented approach was systematically evaluated in our experiments. While this approach introduces a dependency of the training step on the quality of the reference tractogram, we perform further experiments (2) with simulated datasets and the corresponding known ground truth tracts for training. Multiple variations of training and test data are explored to analyze this aspect of our method.

#### Classifier output

The classifier produces a probability *P*(*v*_*i*_) for each of 100 different possible fiber directions *v*_*i*_ (1 ≤ *i* ≤ 100) equally distributed over the hemisphere as well as a non-fiber probability *P*_*nonfib*_.

### 2.2. Streamline progression using neighborhood sampling

At each step of the streamline progression the signal is sampled at *N* positions *p*^*j*^ (1 ≤ *j* ≤ *N*) located on a half-sphere in front of and in distance *r* of the current streamline position *p*. At the seed points, where no previous streamline direction *v*_*old*_ is available, the 2*N* sampling points are distributed over the complete sphere.

Classification is performed at each *p*^*j*^ to infer a weighted local direction proposal *v*^*j*^. The subsequent streamline direction *v* is determined as the normalized sum of the weighted proposals: 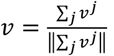 Each proposal direction *v*^*j*^ is determined based on the normalized previous streamline direction *v*_*old*_ probabilities *P*^*j*^(*v*_*i*_) of each possible direction: 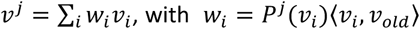. The dot product is a directional prior that promotes straight fibers. An additional hard curvature threshold is employed that, when exceeded, sets *w*_*i*_=0.

If the non-fiber probability of sample *j* exceeds the cumulated weighted probabilities of all possible directions 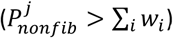, a potential tract boundary is identified and a vote for termination of sample *j* is considered. Now, the position vector *d = p^j^—p* is related to the previous direction *v*_*old*_ in order to decide whether termination is preferable or should be avoided. A termination is considered more likely if non-fiber regions lie straight ahead (i.e. in the current direction of streamline progression *v*_*old*_). If the streamline progresses more or less parallel to the detected fiber bundle margin, a premature termination is rather avoided. To this end an auxiliary sample position *p̂*^*j*^ is evaluated that is determined by a 180° rotation of *d* around *v*_*old*_:

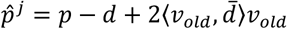

with 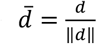 If *P*_*nonfib*_ at the new position *p̂*^*j*^ is > Σ_*i*_*w*_*i*_, *v*^*j*^ is set to (0,0,0) (vote for termination, cf. Figure 1a), otherwise *v*^*j*^ is set to *d̂*=*p̂*^*j*^−*p* to deflect the streamline away from the detected fiber margin (cf. Fig. Figure 1b). A streamline terminates if the majority of all frontal sampling points, meaning sampling points that are located in a 90° cone around *v*_*old*_ ahead of *p,* vote for termination. If no *v*_*old*_ is available, which is the case at the seed location of each streamline, all sampling points have to vote for termination.

**Figure 1:**
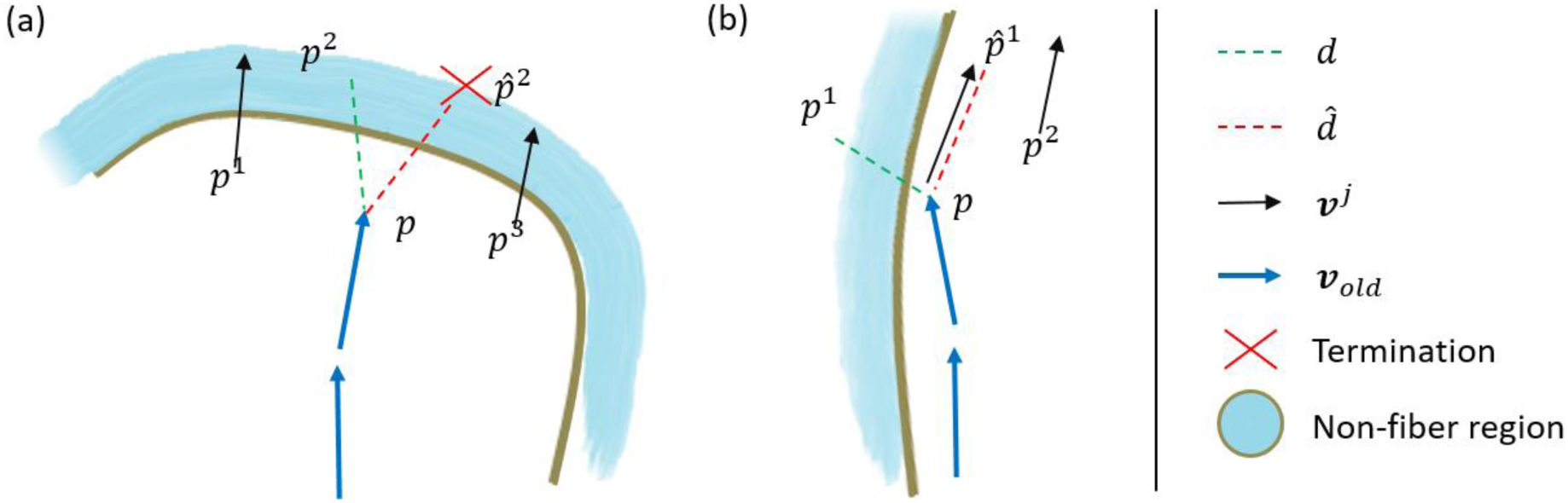
Illustration of the voting process leading to a termination after the next step (a) or to a streamline deflection (b). The current streamline position is denoted *p*. *p*^*j*^ and *p̂*^*j*^ are the sampling points and corresponding alternative sampling points, is the direction proposal at sampling point *p*^*j*^ and *v*_*old*_ is the direction of the previous streamline progression step.

### 2.3. Experiments

We performed four types of experiments to evaluate our approach.

#### Experiment 1

To determine the optimal choice of the tractography algorithm used to create the training data for our approach and to obtain an initial evaluation of its performance, we used a simulated replication of the *FiberCup* phantom (cf. Figure 2) (Fillard et al., 2011). The dataset was simulated using the *Fiberfox* simulation tool (Neher et al., 2014) with the following parameters: 30 gradient directions, a b-value 1000 *s mm*^−2^, 3 *mm* isotropic voxels and a signal-to-noise ratio of about 40. Tractography was performed with 12 combinations of the following openly available tractography and local modelling techniques (cf. Table 1 for corresponding toolkits):

- Tractography algorithms:
  - Deterministic streamline tractography (DET)
  - Fiber assignment by continuous tracking (FACT)
  - Tensor deflection tractography (TEND)
  - Probabilistic streamline tractography (PROB)
  - Global Gibbs tractography

- Local modeling techniques:
  - Single-Tensor model (DT)
  - Two-Tensor model (DT-2)
  - Constrained spherical deconvolution (CSD)
  - Constant solid angle Q-ball (CSA)

**Figure 2:**
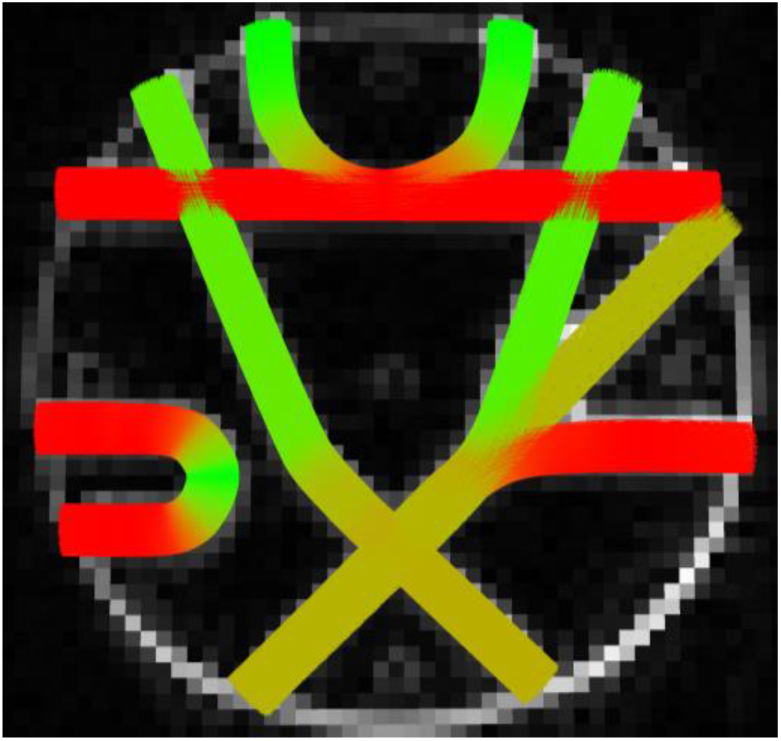
Structure of the *FiberCup* phantom.

**Table 1:** Results of *Experiment 1.* The best scores per metric are highlighted **bold**. The toolkits used to obtain the 12 reference tractograms are ^1^MITK Diffusion (www.mitk.org/wiki/DiffusionImaging), ^2^Camino (camino.cs.ucl.ac.uk/) and ^3^MRtrix (www.mrtrix.org, v0.2).

| Model | Type | NC | VC | IC | OL | AE |
| --- | --- | --- | --- | --- | --- | --- |
| DT<sup>1</sup> | DET<sup>1</sup> | 60% | 25% | 15% | 21% | 5° |
| DT<sup>1</sup> | FACT<sup>1</sup> | 62% | 23% | 14% | 24% | 6° |
| DT<sup>1</sup> | TEND<sup>1</sup> | 84% | 8% | 8% | 21% | 10° |
| DT<sup>2</sup> | PROB<sup>2</sup> | 57% | 23% | 20% | 27% | 8° |
| DT<sup>1</sup> | Global<sup>1</sup> | 82% | 10% | 8% | 42% | 12° |
| CSA<sup>1</sup> | DET<sup>3</sup> | 24% | 67% | 9% | 70% | 8° |
| CSA<sup>1</sup> | PROB<sup>3</sup> | 91% | 5% | 4% | 83% | 18° |
| CSA<sup>1</sup> | Global<sup>1</sup> | 81% | 14% | 5% | 74% | 13° |
| DT-2<sup>2</sup> | DET<sup>2</sup> | 60% | 37% | 3% | 58% | 6° |
| CSD<sup>3</sup> | DET<sup>3</sup> | 21% | 78% | **1%** | 86% | **4°** |
| CSD<sup>3</sup> | PROB<sup>3</sup> | 66% | 28% | 7% | 93% | **4°** |
| CSD<sup>3</sup> | Global<sup>1</sup> | 81% | 17% | 2% | 72% | 12° |
| - | Proposed | **3%** | **93%** | 4% | **94%** | **4°** |

The presented approach was trained individually with each of the 12 benchmark methods. Based on the results of this analysis, the most promising algorithm was chosen to obtain the training tractogram for all further experiments (*A*_*train*_).

To quantify the performance of the methods, the following metrics from the *Tractometer* evaluation protocol (Côté et al., 2013) were analyzed: the fraction of no connections (NC), valid connections (VC), invalid connections (IC) and bundle overlap (OL), as well as an additional measure for the local angular error (AE) (Neher et al., 2015a).

Seeding was performed homogeneously within the image. Not white-matter mask was used to constrain the tractography. All tractography algorithms were run with their default parametrization. Only the stopping criteria (FA and ODF peak thresholds) were manually adjusted to obtain plausible results (FA threshold 0.15, peak threshold for CSA 0.085 and peak threshold for CSD 0.15).

The presented method was run with *N*=50 sampling points, a step size of 0.5⋅*f* (*f* is the minimal voxel size in *mm*), *r*=0.25⋅*f*, a minimum fiber length of 20*mm*, a maximum fiber length of 200*mm*, and a hard curvature threshold at a maximum angle of 45° between two steps or a maximum directional standard deviation of 30° over the last centimeter. The classifier was trained using 30 trees, a maximum tree depth of 50 and a Gini splitting criterion. The training data was sampled equidistantly (0.5⋅*f*) along the input tractogram fibers as well as on 50 randomly placed points in each non-fiber voxel. The defaults for step size and angular thresholds were empirically determined and are typical for streamline based fiber tractography approaches. The number of sampling points and the forest parameters yielded stable results in a broad range and increasing them further would mainly impact the computational cost of the method. In this experiment we used the raw diffusion-weighted signal values, resampled to 100 directions equally distributed over the hemisphere using spherical harmonics (cf. sec. 2.1), as input for the classifier.

#### Experiment 2

The *in vivo* performance of our approach in comparison to the 12 methods described in *Experiment 1* was qualitatively evaluated on basis of reconstructions of the corticospinal tract (CST) and by an analysis of the spatial distribution of fiber end points. The dataset was acquired using 81 gradient directions, a b-value 3000 *s mm*^−2^ and 2.5 *mm* isotropic voxels. All methods were run with their default parameterization, which is the same as in *Experiment 1*. In contrast to *Experiment 1*, no manual adjustment of FA and ODF peak thresholds was necessary *in vivo*. As in *Experiment 1*, the interpolated raw signal values were used as input features for the classifier. To obtain the training reference, we used the method determined in *Experiment 1* (*A*_*train*_).

#### Experiment 3

In this experiment, we assessed the performance of the presented method using the ISMRM tractography challenge 2015 data (Maier-Hein et al., 2016) (www.tractometer.org/ismrm_2015_challenge/). The ground truth fiber bundles mimic the shape and complexity of 25 well known *in vivo* fiber bundles (cf. Figure 3). The diffusion-weighted dataset was simulated with 32 gradient directions, a b-value 1000 *s mm*^−2^ and 2 *mm* isotropic voxels.

This experiment enabled us to compare our approach to all 96 original submissions of the tractography challenge comprising a large variety of tractography pipelines with different pre‐processing, local reconstruction, tractography and post‐processing algorithms.

As preprocessing step, the dataset was denoised and corrected for distortions using *MRtrix* (*dwidenoise* & *dwipreproc*, http://www.mrtrix.org/).

The random forest classifier was trained using all combinations of the following parameters, resulting in a total of 8 trained classifiers:

- Classification features:
  1. Interpolated raw signal values
  2. Spherical harmonics coefficients (order 6)
- Additional features:
  1. no additional features
  2. T1 signal and GFA
- Training tractograms:
  1. Ground truth fibers (*GT*_*train*_) used to simulate the phantom image
  2. Tractogram obtained using the method *A*_*train*_

To reduce the computational load and since the parameters yielded stable results across a broad range, the maximum tree depth was reduced to 25 and the number of sampling points *N* during tracking was reduced to 30. To obtain balanced classes, the number of non-fiber samples was chosen automatically to match the number of fiber samples.

For each of the 8 classifiers, tractography was performed two times using 1 and 3 seed points per voxel, respectively.

The same evaluation metrics as presented in the original challenge were analyzed for all 16 tractograms using the official evaluation pipeline: valid bundles (VB), invalid bundles (IB), valid connections (VC), bundle overlap (OL) and bundle overreach (OR).

**Figure 3:**
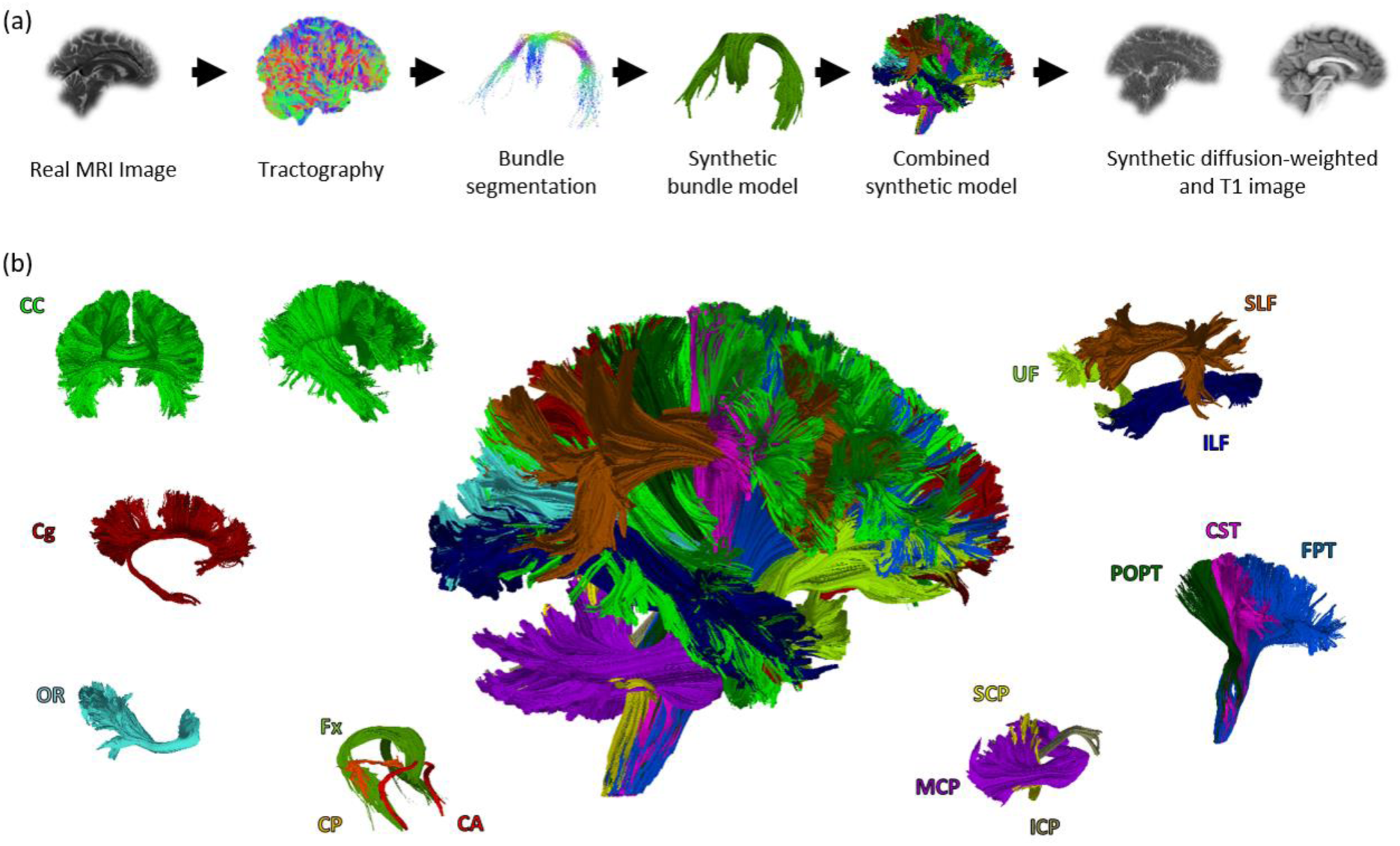
Illustration of the phantom generation process (a) and its constituent 25 fiber bundles (b). For details about the ISMRM tractography challenge 2015 and the phantom, please refer to (Maier-Hein et al., 2016) and the challenge homepage www.tractometer.org/ismrm_2015_challenge/.

#### Experiment 4

Our final experiments aim at evaluating the generalization capability of the presented method using *in vivo* and *in silico* data:

As *in vivo* data we used five datasets of the Human Connectome Project (HCP) (Van Essen et al., 2012, 2013; Van Essen and Ugurbil, 2012) for training and five other HCP datasets for testing (*HCPtrain* [Subject IDs: 984472, 979984, 978578, 994273, 987983] and *HCPtest* [Subject IDs: 992774, 991267, 983773, 965771, 965367]). The HCP datasets are acquired using 270 gradient directions, three b-values (1000 *s mm*^−2^, 2000 *s mm*^−2^, 3000 *s mm*^−2^) and 1.25 *mm* isotropic voxels.

As *in silico* data for this experiment, we employed the IMSRM tractography challenge phantom already used in *Experiment 3*. Since the phantom and the HCP datasets were acquired with different imaging sequences, they feature different image contrasts. Therefore, the phantom dataset was normalized to feature the same mean and standard deviation inside the white matter as the HCP datasets.

Three types of generalization were tested:

1. *In vivo* → *in vivo*: In this part of the experiment, we trained our method using *HCP*_*train*_ and evaluated its performance on the unseen datasets of *HCP*_*test*_. Training was performed using multi-tissue CSD (Jeurissen et al., 2014) deterministic tractography as *A*_*train*_, about 12 million samples per dataset and spherical harmonics coefficients as classification features calculated from the *b* = 3000 *s mm*^−2^ shell of *HCP*_*train*_. We only used the highest b-shell instead of all three available shells to avoid to triple the amount of required memory for the features. Since the HCP datasets feature a much higher resolution compared to the datasets used in the other experiments, tractography on *HCP*_*test*_ was performed using an increased sampling distance (0.7⋅*f*). Streamlines were seeded five times in every brain voxel. Evaluation was performed qualitatively by manually extracting the corticospinal tract (CST), cingulum (Cg) and fornix (Fx) from each of the five test results and a successive visual inspection of the tracts.
2. *In vivo* → *in silico*: In this part of the experiment, we trained our method using *HCP*_*train*_ and evaluated its performance on the unseen phantom dataset already used in *Experiment 3*. The same tractography parameterization as in *Experiment 3* and the same training tractograms already employed in part (1) were used. To match the b-value of the phantom dataset, the spherical harmonics coefficients from the *b* = 1000 *s mm*^−2^ shell of *HCP*_*train*_ were used as training features. The resulting tractogram was evaluated quantitatively using the same measures already described in *Experiment 3*.
3. *In silico* → *in vivo*: In this part of the experiment, we trained our method on the phantom dataset and the corresponding ground truth fibers introduced in *Experiment 3* and tested it on the five *in vivo HCP*_*test*_ datasets. As in (2), the *b* = 1000 *s mm*^−2^ shells of *HCP*_*test*_ datasets were used to calculate the features for our method. Evaluation and tractography was performed analogous to (1).

## 3. Results

### Experiment 1

An overview The best results on the phantom image were obtained using the CSD DET tractography (Tournier et al., 2007, 2012) for training our approach (*A*_*train*_ = CSD DET). With this configuration, the presented approach outperformed all benchmark methods in four out of the five metrics (cf. Table 1). Only 3% of the tracts terminated prematurely. Furthermore, the presented approach yielded the highest percentage of valid connections (93%), the highest bundle overlap (94%) and the lowest local angular error (4%). All 7 valid bundles in the phantom were reconstructed successfully. Also, the percentage of invalid connections (4%) is rather low compared to the majority of benchmark algorithms (rank 4 out of 13). When varying the method that was used for training, the percentage of prematurely ending fibers and valid connections yielded by the presented approach improved on average by 56% and 36% respectively as compared to the benchmark tractograms. The average percentage of invalid connections, however, was increased by 21%.

### Experiment 2

Based on the results of *Experiment 1*, the CSD DET tractography was used as training method *A*_*train*_ for the *in vivo* experiments. *In vivo*, our approach successfully reconstructed a whole brain tractogram including challenging regions such as the crossing between the corpus callosum, the CST and the superior longitudinal fasciculus. Our method was furthermore able to reconstruct parts of the CST that other approaches often missed (cf. lateral projections of the CST in Figure 4b). In comparison to the benchmark algorithms, most of the fibers reconstructed by the presented approach correctly terminated in the cortex (cf. Figure 4a).

**Figure 4:**
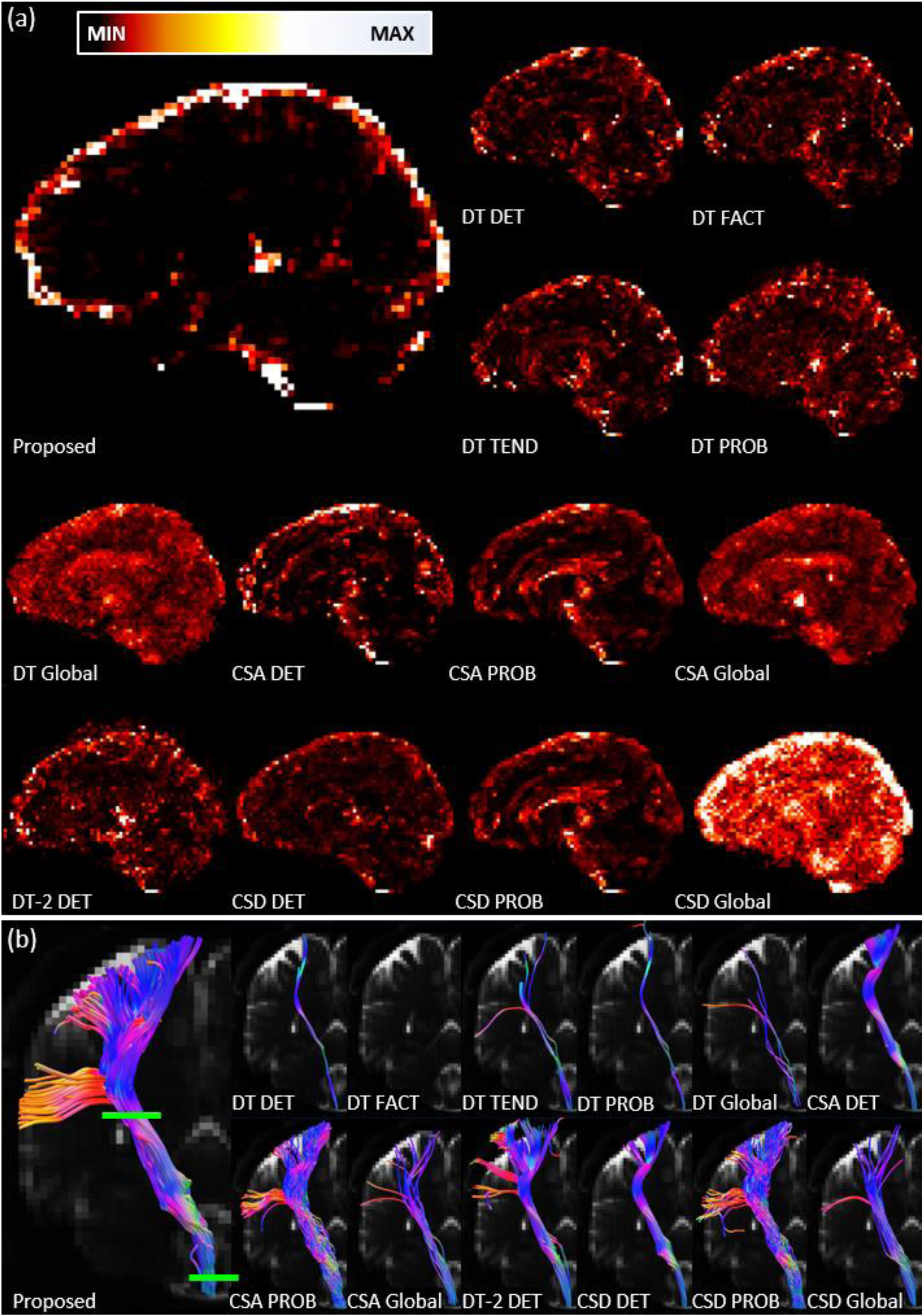
Results on the *in vivo Experiment 2.* (a) shows the max-normalized voxel-wise number of fiber endpoints, maximum intensity projected over 20 sagittal slices. (b) shows the corticospinal tracts obtained with all 13 algorithms. The green bars schematically depict the inclusion regions used for all whole brain tractograms to extract the respective CST.

### Experiment 3

Figure 5 depicts the results of our approach in comparison to the original challenge results. For reference, the same numbering of the teams as in the challenge paper were used. Team number 1-20 belong to the 20 teams that submitted results to the challenge. To obtain a clear presentation of our results, we split them into two groups (Team 21 and 22) according to the respectively employed training tractogram (Tractogram obtained with *A*_*train*_ or the ground truth tracts *GT*_*train*_). In the following paragraphs we analyze our results in comparison to the original tractography submissions and with respect to the different training and tractography parameterizations described in Section 2.3, *Experiment 3*.

#### General observations

The presented approach is the only method that was able to reconstruct all 25 valid bundles. Regardless of the pipeline configuration, the minimum number of valid bundles reconstructed by our approach was 24, which could not be matched by any other approach. Furthermore, none of the original 96 submissions reached valid connection and bundle overlap ratios larger than 50% while keeping the overreach ratio smaller than 50%, in contrast to all 16 new tractograms. Compared to the mean scores over all 96 benchmark submissions, the mean scores over the new tractograms were improved for all metrics except the bundle overreach (VB: +2.82, IB: -1.49, VC: +14.83, OL: +31.26, OR: +12.09). While the overreach was increased by 12.09%, the increase in bundle overlap was almost three times as high.

#### Training data (Team 21 vs. 22)

As expected, the presented method performs clearly better when using the *GT*_*train*_ tracts as training data as compared to the training tractogram obtained with *A*_*train*_. Especially the valid connections, bundle overlap and bundle overreach are distinctly improved using the ground truth.

#### Effect of the different diffusion-weighted features

Using spherical harmonics coefficients instead of the raw signal values as classification features results in consistently higher valid connection ratios and a higher overlap at the cost of a lower valid bundle score (maximum VB is 24) and a slightly higher overreach.

#### Effect of additional features

The differences are very small and consistent effects could only detected when training on the ground truth (Team 22), where inclusion of T1 and GFA features lead to a slight increase of valid connections, a higher overlap and a lower overreach.

#### Effect of the varying number of seed points

Using a larger number of seed points resulted in a higher sensitivity – all configurations that yielded 25 valid bundles used 3 seed points – and a higher bundle overlap, while at the same time increasing the overreach and number of invalid bundles.

**Figure 5:**
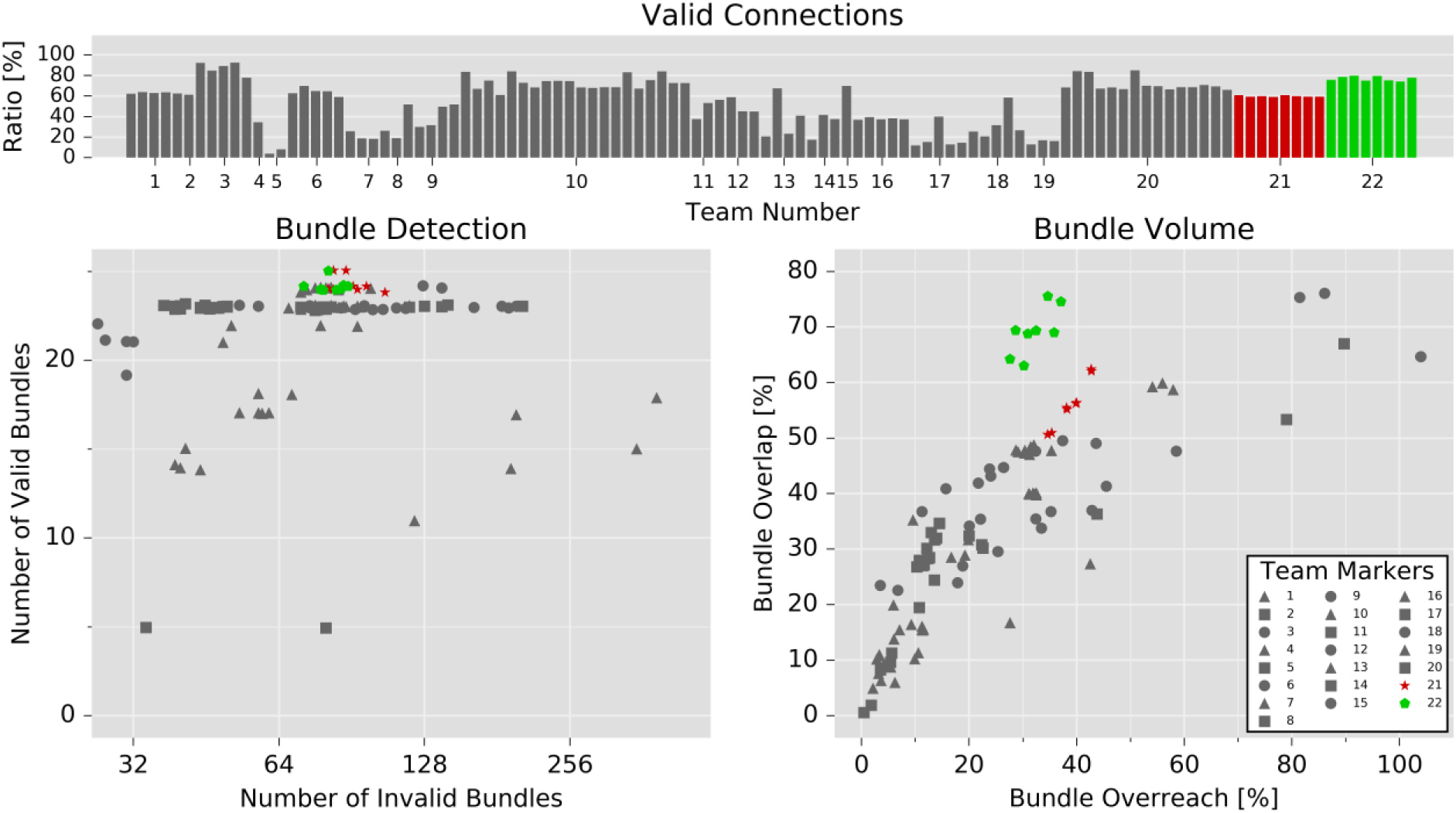
Scores of the 16 new tractograms obtained using the presented method (color) in comparison to the original challenge submissions (gray). The tractograms in Team 21 (red, star) were obtained using *A*_*train*_ to generate the training reference and the tractograms in Team 22 (green, pentagon) with the ground truth tracts *GT*_*train*_ as training reference.

### Experiment 4

Overall, the presented approach was able to generalize well to unseen datasets. The following paragraphs describe the individual results of the three parts of this experiment.

1. *In vivo* → *in vivo*: Figure 6 shows the tracts extracted from the five HCP test tractograms. All tracts were reconstructed successfully, which demonstrates that the classification based approach is capable of generalization to unseen datasets.
2. *In vivo* → *in silico*: The method trained on five HCP datasets was able to reconstruct 24/25 valid bundles in the ISMRM tractography challenge phantom, while reconstructing 88 invalid bundles, which is comparable to the results of *Experiment 3*, where the approach was directly trained on the phantom dataset. The fraction of valid connections (38%) and the bundle overlap (46%) on the other hand were considerably lower. With 34%, the bundle overreach was again similar to the results of *Experiment 3*.
3. *In silico* → *in vivo*: As in (1), the approach trained on simulated data was able to reconstruct the 3 tracts of interest in all five unseen *in vivo* test datasets (cf. Figure 7).

**Figure 6:**
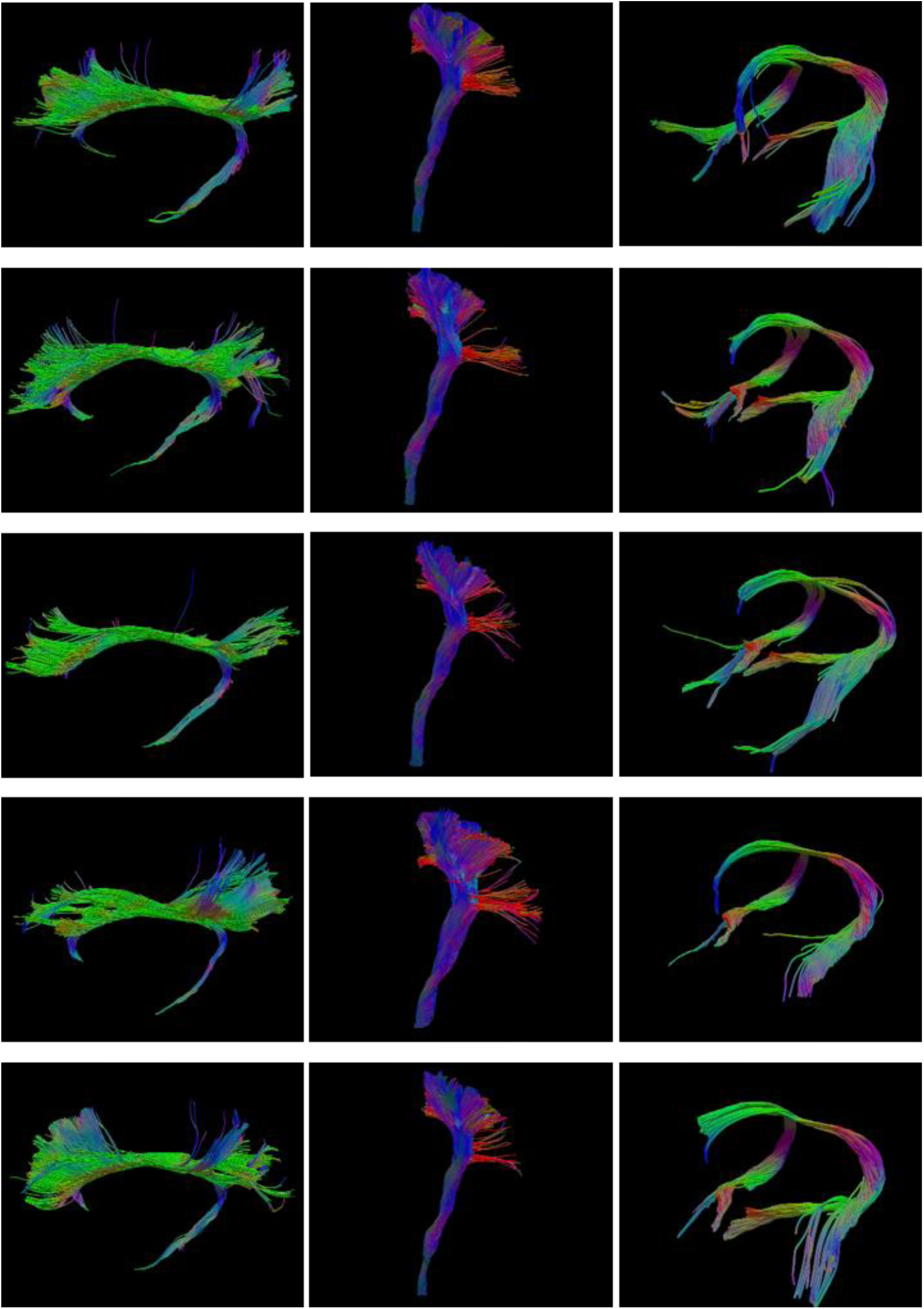
Cingulum (left), corticospinal tract (middle) and fornix (right) reconstructed from the five HCP test subjects (cf. sec. 2.3 *Experiment 4 (1)).* The results were obtained with the presented approach after training on five different HCP subjects.

**Figure 7:**
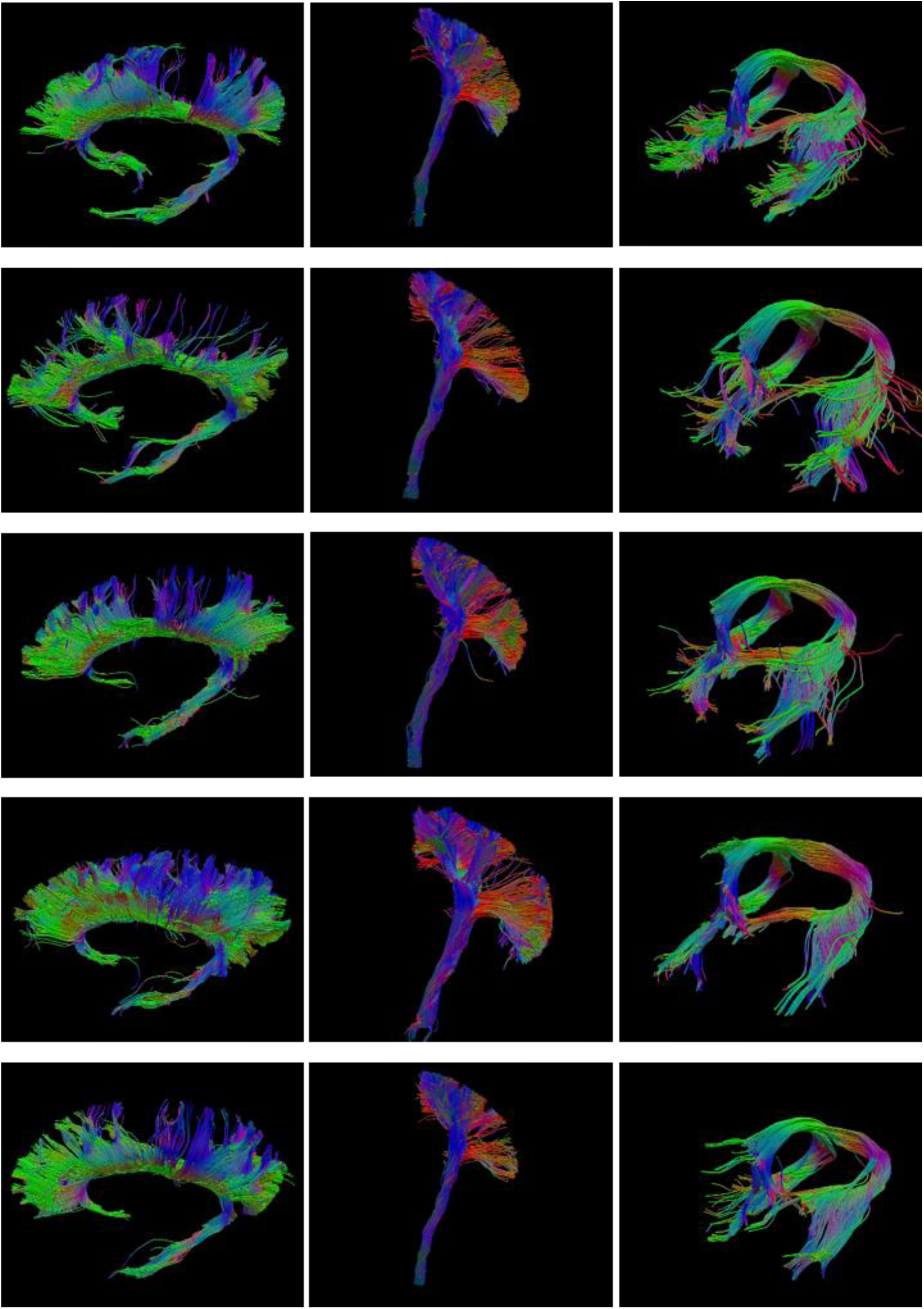
Cingulum (left), corticospinal tract (middle) and fornix (right) reconstructed from the five HCP test subjects (cf. sec. 2.3 *Experiment 4 (3)).* The results were obtained with the presented approach after training on the ISMRM tractography challenge phantom.

## 4 Discussion and Conclusion

We presented a random-forest classification-based approach to fiber tractography using neighborhood information that guides each step of the streamline progression.

The presented approach is the first to utilize machine learning for fiber tractography. The method systematically exploits the diffusion-weighted signal not only locally but also in the neighborhood of the current streamline position in order to increase sensitivity, which is rarely done in current tractography pipelines.

We thoroughly evaluated the performance of the presented method in comparison to over 100 state-of-the-art tractography pipelines on simulated phantom datasets as well as *in vivo*.

In the *in vivo* experiments (*Experiment 2*), our approach yielded very good results in reconstructing difficult tracts (e.g. the lateral projections of the CST) and a much lower number of fibers ending prematurely inside the brain. As expected, tensor based approaches had difficulties in detecting the lateral projections of the CST. However, even the benchmark method that performed best in the phantom experiments (*A*_*train*_ = CSD DET) was unable to detect these projection fibers. The benchmark methods that showed a relatively high sensitivity in this region (CSA+CSD PROB and DT-2 DET) displayed a very low specificity in the phantom experiments (*Experiment 1*) as well as with respect to the *in vivo* end-point distribution. In contrast, the presented algorithm showed a constantly high sensitivity and specificity.

In the quantitative analysis on the ISMRM tractography challenge phantom dataset (*Experiment 3*), our approach showed the highest sensitivities in terms of reconstructed valid bundles while still yielding a specificity in terms of reconstructed invalid bundles above average. It is furthermore the only approach to reach bundle overlap scores above 50% while at the same time keeping the bundle overreach below 50%.

The capability of the presented approach to generalize to unseen datasets was successfully demonstrated in *Experiment 4*. We showed that generalization is possible between different *in vivo* images acquired with the same MR sequence as well as between *in vivo* images and simulated datasets (in both directions). The latter experiment, generalizing from simulated datasets to *in vivo*, is especially important, since it enables us to train on datasets with known ground truth. This eliminates the bias introduced by training on tracts obtained with a conventional tractography method.

While the presented results are promising, there are still some challenges to address. Further work is necessary to quantify and improve the performance of the presented approach when training on simulated datasets. One important aspects in this regard is the simulation of further datasets to obtain more comprehensive training data with respect to image contrast and fiber structure. Another aspect where methodological improvements are definitely possible is the currently employed naïve approach to generalize between the simulated and *in vivo* domain, using approaches of unsupervised domain adaptation and transfer learning (Götz, Michael et al., 2014; Heimann et al., 2014; Long et al., 2016, 2014; McKeough et al., 2013; Pan and Yang, 2010; Sener et al., 2016). Another challenge is the still high number of invalid bundles, which is a known issue of current fiber tractography approaches (Maier-Hein et al., 2016). This is a challenge the whole fiber tractography community is facing and that does not have a simple solution. However, it seems promising to incorporate as much additional knowledge in the tractography process as possible, for which machine learning based approaches seem to be well suited. Interesting candidates would be functional MRI data or prior knowledge in form of cortical parcellations. An extension of the presented method to directly include a distinction between different non-white matter tissue types, such as gray matter and corticospinal fluid, seems promising to further improve the decision on where to terminate the fiber progression. This includes addressing further interesting aspects of the fiber termination such as orthogonality to and uniform coverage of the gray-white-matter interface. We are also planning to analyze how the process of including neighborhood information can be improved further, e.g. by applying patch-based classification at the current streamline position in order to obtain a more ‘global’ view on the image that might be necessary to avoid invalid connections.

The source-code of all methods presented in this work is available open-source and integrated into the Medical Imaging Interaction Toolkit (MITK) (Fritzsche et al., 2012; Nolden et al., 2013). The datasets used in *Experiment 1* and *2* are available for download at www.nitrc.org/projects/diffusion-data/. The dataset used in *Experiment 3* as well as many other resources regarding the ISMRM tractography challenge are available at www.tractometer.org/ismrm_2015_challenge/. The HCP datasets used in *Experiment 4* are available on www.humanconnectome.org/data/.

## Acknowledgments

Data were provided in part by the Human Connectome Project, WU-Minn Consortium (Principal Investigators: David Van Essen and Kamil Ugurbil; 1U54MH091657) funded by the 16 NIH Institutes and Centers that support the NIH Blueprint for Neuroscience Research; and by the McDonnell Center for Systems Neuroscience at Washington University.

This work was supported by the Collaborative Research Center (SFB/TRR 125 Cognition-Guided Surgery) of the German Research Foundation (DFG) grant number INST 35/1120-1, DFG grant MA 6340/10-1 and DFG grant MA 6340/12-1.

